# A new fluorescence-based approach for direct visualization of coat formation during sporulation in *Bacillus cereus*

**DOI:** 10.1101/2023.05.12.540479

**Authors:** Armand Lablaine, Stéphanie Chamot, Mónica Serrano, Cyrille Billaudeau, Isabelle Bornard, Rut Carballido-Lopez, Frédéric Carlin, Adriano O. Henriques, Véronique Broussolle

**Affiliations:** INRAE, Avignon Université, UMR SQPOV, F-84000 Avignon, France; Instituto de Tecnologia Quimica e Biologica, Universidade Nova de Lisboa, 2780-157 Oeiras, Portugal; MICALIS Institute, INRAE, AgroParisTech, Université Paris-Saclay, 78350 Jouy en Josas, France; INRAE, Pathologie végétale, F-84143 Montfavet, France

## Abstract

The pathogenic bacteria *Bacillus cereus*, *Bacillus anthracis* and *Bacillus thuringiensis* form spores encased in a protein coat surrounded by a balloon-like exosporium. These structures mediate spore interactions with its environment, including the host immune system, control the transit of molecules that trigger germination and thus are essential for the spore life cycle. Formation of the coat and exosporium has been traditionally visualized by transmission electronic microscopy on fixed cells. Recently, we showed that assembly of the exosporium can be directly observed in live *B. cereus* cells by super resolution-structured illumination microscopy (SR-SIM) using the membrane MitoTrackerGreen (MTG) dye. Here, we demonstrate that the different steps of coat formation can also be visualized by SR-SIM using MTG and SNAP-cell TMR-star dyes during *B. cereus* sporulation. We used these markers to characterize a subpopulation of engulfment-defective *B. cereus* cells that develops at a suboptimal sporulation temperature. Importantly, we predicted and confirmed that synthesis and accumulation of coat material, as well as synthesis of the σ^K^-dependent protein BxpB, occur in cells arrested during engulfment. These results suggest that, unlike the well-studied model organism *Bacillus subtilis*, the activity of σ^K^ is not strictly linked to the state of forespore development in *B. cereus*.

## Introduction

Sporulation is a developmental process in which a vegetative cell transforms into a highly resilient spore (Fig. 1). Among the spore forming bacteria, endospores (hereinafter simply referred to as ‘spores’) made by *Firmicutes* species, which include the model organism for developmental studies *Bacillus subtilis*, the foodborne pathogen *B. cereus*, its closely relatives *B. anthracis* and *B. thuringiensis,* and the nosocomial pathogen *Clostridioides difficile*, are of particular interest for developmental biology or clinical studies^1, 2^.

**Figure 1.**
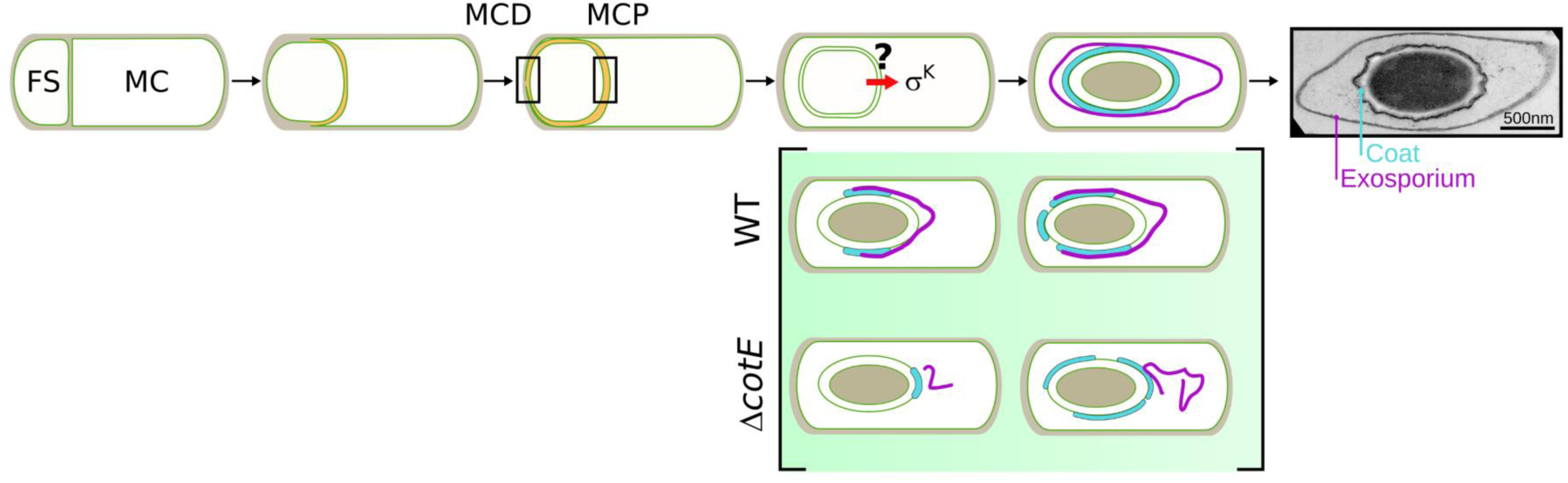
Assembly of proteinaceous spore surface layers during *Bacillus cereus* sporulation. Schematic illustration of *B. cereus* sporulation stages and transmission electron microscopy (TEM) image of a *B. cereus* ATCC 10876 spore. A tight control coordinates σ^K^ activation and engulfment completion across *Firmicutes* species, but whether σ^K^ activation strictly depends on engulfment completion in *B. cereus* is unknown. Sequence of coat (cyan) and exosporium (pink) formation as described by TEM in wild-type (WT) and Δ*cotE* strains^15, 18^. Similar phenotypes were described in *B. anthracis*. In absence of exosporium cap in Δ*cotE* mutant sporangia, coat deposition sequence is modified compared to that of the WT. MCD, Mother Cell Distal; MCP, Mother Cell Proximal pole of the forespore; FS, Forespore; MC, Mother Cell.

Transmission electronic microscopy (TEM) has traditionally been the technique used to study the formation and architecture of *Firmicutes* spores^3^. Pioneer TEM studies revealed that the general architecture of spores and the sporulation steps are conserved among *Firmicutes* species^4^. Firstly, the cell entering sporulation divides asymmetrically into a small cell, the forespore and a larger mother cell, both defining a sporangium (Fig. 1). Then, the mother cell membrane migrates around the forespore in a phagocytosis-like process so-called engulfment. Ultimately, the mother cell lyses releasing a mature spore in the environment (Fig. 1). The mature spore consists in a series of concentric shells, with from the center to the outside, the dehydrated core containing the genetic information, the primordial germ cell wall, the cortex, a layer of modified peptidoglycan, which is wrapped in a complex proteinaceous external shell. The architecture of the proteinaceous surface shell differs greatly between bacterial species^2, 5, 6^, likely reflecting the niches in which spores persist and eventually germinate, ranging from the soil, the oxygen-deprived mammalian gut, or the inhospitable macrophages^2, 7^. Hence, while the spores of *B. subtilis* are encased in a multilayered thick proteinaceous coat, the spores of species belonging to the *B. cereus* group are enclosed in a thin layer of coat surrounded by a balloon-like exosporium, both structures being separated by an interspace presumably filled by polysaccharides (Fig. 1)^8^.

Compared to other *Firmicutes* spore-formers, and thanks to its high natural transformability, the assembly of the multilayered coat of *B. subtilis* has been studied in depth and is now well understood ^1, 6, 9, 10^. Notably, most of the known coat proteins of *B. subtilis* were localized during sporulation by conventional fluorescence microscopy, using GFP-fusions inserted at the original genetic locus and the model of *B. subtilis* coat assembly largely originates from those studies^1, 6, 9^. In contrast, for species of the *B. cereus* group, genetic manipulations remain limited by its low transformability^11, 12^ and GFP or mCherry fusions are, most of the time, carried on shuttle plasmids^13–21^. In addition, for *B. cereus sensu stricto* species, and in contrast to *B. anthracis,* GFP-fusions were ineffective to localize early or late assembling exosporium proteins^13, 22^. Hence, the model of coat and exosporium assembly mainly originates from TEM analysis of fixed cells (Fig. 1)^15, 18, 23–26^. Coat assembly appears to differ greatly in *B. subtilis* and *B. cereus,* as the initiation of this assembly process occurs in opposite regions of the forespore^15, 18^. Despite such a fundamental difference, formation of the *B. cereus* coat remains a poorly studied process and an alternative way allowing the direct observation of proteinaceous layers formation in live cells would be of interest to better characterize this developmental process.

Although coat assembly begins early after formation of the mother cell among *Firmicutes* species, the coat is clearly visible by classical TEM only after engulfment completion and is not detected in Δ*sigK* sporangia^5, 15, 18, 23, 24, 27–29^. In *B. subtilis*, activation of the late sporulation sigma factor σ^K^ is tightly coupled to engulfment completion through several levels of regulation, including delayed transcription of the *sigK* gene, removal of an inhibitory pro−sequence and excision of a prophage-like skin element from the σ^K^-encoding gene^29–33^. Despite a high conservation of sigma factor primary sequence, difference in the control of SigK activation had been reported among *Firmicutes* species^29^. Notably, in *C. difficile*, an inhibitory σ^K^ prosequence is absent and σ^K^ activity occurs during engulfment as well as in engulfment-defective cells^34–38^. The activity of σ^K^ in engulfment-defective cells leads to the synthesis and the accumulation of electron-dense coat material at, or close to the forespore surface, observed by TEM and referred as a “coat bearding” phenotype^35^. For most *B. cereus* strains, a σ^K^ pro-sequence is present, while the *sigK* gene is not interrupted by a prophage sequence as in *B. subtilis* and *C. difficile*. Whether σ^K^ activation and accumulation of coat material is linked to engulfment completion in *B. cereus* is not known.

We recently showed that the exosporium layer could be visualized in *B. cereus* live sporulating cells and spores using the membrane dye Mitotracker green (MTG) and Structured Illumination Microscopy (SIM)^15^. Using MTG and a SNAP-fusion for the exosporium cap protein CotY, we showed the existence of a second small exosporium cap at the mother cell proximal pole (MCD pole) of Δ*exsY* spores not previously observed by TEM ^25, 26^. This indicates that exosporium formation shares more similarities with the *B. subtilis* model of coat formation than expected^1, 2, 39, 40^. Here, we show that SNAP-cell TMR-star (TMR-star), a fluorescent substrate used to label SNAP-tag fusion proteins, and MTG are both able to specifically bind to the coat surface of *B. cereus*. The resulting fluorescent signals observed in sporangia and in spores can be used to determine precisely the stage of coat assembly. Using SIM analysis, we observe fluorescence signals that we assigned to an accumulation of coat material in engulfment-defective *B. cereus* cells, which was confirmed by TEM analysis. Finally, we demonstrate that the SNAP-tag fusion allows to successfully detect the late assembling BxpB exosporium protein and that both the TMR-star association with the *B. cereus* coat and the TMR-star signal labeling the BxpB-SNAP fusion are detected in engulfment-defective *B. cereus* cells. Our results provide a new way to evaluate the state of surface layers formation in live sporangia and suggest that regulation of σ^K^ activation is weakly linked to the state of forespore development in the *B. cereus* group, indicating that pathogenic spore-formers deploy different strategies to control this step of spore formation.

## Results

### TMR-star and MTG bind to the forespore surface and their signal localization reflects the pattern of coat deposition

We previously used the self-labelling SNAP-tag to determine the localization of early assembling exosporium proteins during *B. cereus* sporulation^15^. Indeed, SNAP-protein fusions can be localized by fluorescence microscopy upon addition of a cell-permeable fluorescent ligand^41^. Surprisingly, we noticed that the TMR-star, a red fluorescent SNAP substrate, also labeled the surface of spores that do not harbor any SNAP-tag fusion (Fig. 2A, TMR channel in panel *h*)^15^. The coat of *Firmicutes* spores bind different fluorescent substrates^37,42–46and^ in *B. cereus* ATCC 14579 wild-type cells (WT14579), the TMR-star signal displayed a particular ellipsoid shape (Fig. 2A, panel *h*, cyan arrows) inside the exosporium layer (Fig. 2A, panel *h*, pink arrow), matching the localization of the coat. We thus wondered whether binding of TMR-star to the spore surface of *B. cereus* could be used as a coat marker in fluorescence microscopy studies. Coat deposition in the *B. cereus* group is a sequential process (Fig. 1), with electron-dense coat material firstly appearing on the long side of the forespore^15, 18, 23^. In the absence of the exosporium cap, as in cap-defective Δ*cotE* cells, the sequence of coat deposition is altered and coat deposition can initiate on the forespore poles (Fig. 1)^15, 18^. We reasoned that, if TMR-star binds to *B. cereus* coat, intermediate TMR-star signals should be observed during sporulation and would be impacted by the absence of the exosporium cap. Thus, we carefully examined the TMR-star signal during the sporulation of WT14579 (Fig. 2A) and of a congenic Δ*cotE* mutant (Fig. 2B). We induced sporulation by resuspension in SMB at 20°C, since Δ*cotE* spores can be partly decoated at this suboptimal temperature^47^. Sporulating cells and spores collected throughout sporulation were labeled with TMR-star and the MTG membrane dye and imaged with bright field (BF) and super-resolution SIM illumination modes.

**Figure 2.**
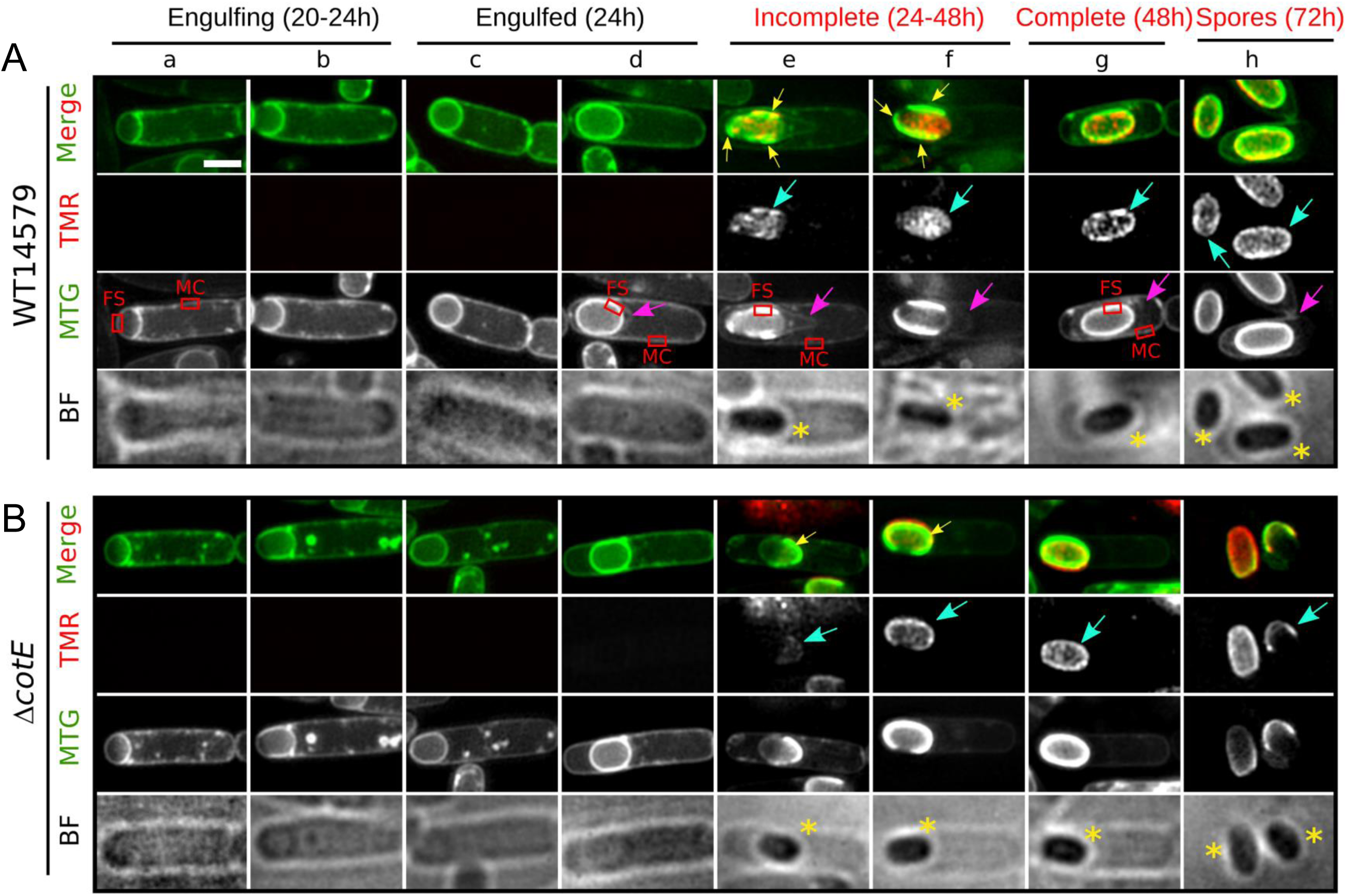
TMR-star and MTG bind to the forespore surface after engulfment completion. (**A**) WT14579 or (**B**) congenic Δ*cotE* cells sporulating at 20°C in SMB were labeled with TMR-star (TMR, red on merged images) and MTG (green on merged images) dyes and imaged with structured illumination microscopy (SIM) and bright field (BF) microscopy. Representative sporangia (panels *a*-*g*) or spores (panel *h*) observed at indicated times after resuspension in SMB medium are shown. Sporulation stages are indicated and stages written in red point the presence of TMR-star and MTG bright signals on the forespore surface. Yellow arrows bright MTG signals reflecting patterns of coat deposition according to previous TEM analysis in similar genetic backgrounds. Cyan arrows point to the TMR-star signal observed on the forespore surface. Yellow asterisks indicate refringent forespores and spores. Pink arrows point to the exosporium. Red boxes in MTG channel illustrate areas used to perform quantification of FS/MC MTG signal intensity ratio, as presented in Fig. 3. Scale bar in panel *a* represents 1 μm.

In both WT14579 and Δ*cotE* sporangia, the TMR-star signal appeared after engulfment and was first observed incompletely covering the surface of engulfed forespore (Fig. 2A and B, panels *e* and *f*, cyan arrows). Under BF illumination, the forespore of those sporangia appeared dark, indicating the development of forespore refringency (panels *e* and *f*, yellow asterisks). In WT14579 sporangia showing a TMR-star signal partially covering the forespore surface (in the MTG channel), the basal layer of the exosporium cap appeared clearly more extended from the forespore membrane (Fig. 2A, panels *e* and *f*, 504±109 nm, n=35), than in WT14579 sporangia without TMR-star signal (panel *d*, 219±73 nm, n=48). This result indicates that binding of TMR-star on the forespore surface is concomitant to exosporium extension. Visual inspection of SIM reconstructed images also revealed a brighter MTG signal partly covering the periphery of refringent forespores with a TMR-star signal (Fig. 2A and B, panels *e-f*). The pattern of MTG brighter signal localization differed between WT14579 and Δ*cotE* sporangia (panels *e* and *f* in Fig. 2A and B) but remarkably, in both cases, this localization fully reflected the patterns of coat deposition previously described by TEM in *B. cereus* and in *B. anthracis* (Fig. 1)^15, 18, 23^. Finally, in 50±7% of Δ*cotE* spores (144/263 cells from two independent experiments), both the MTG and the TMR-star signals partially covered the spore surface (Fig. 1B, panel *h*, see Materials and methods), while a total coverage was observed in WT14579 spores (Fig. 1A, panel *h*). We concluded that the TMR-star and MTG bright signals observed in sporangia and in spores match all features of coat assembly previously described by TEM in both *B. cereus* and *B. anthracis*^15, 18, 23, 47^.

### Quantification of MTG signal intensity in sporangia

To determine if the brighter MTG signals observed on the periphery of refringent forespores reflect MTG binding to additional structures (other than the cell membrane), we quantified the ratio of the MTG signal intensity between the forespore and the mother cell (FS/MC MTG fluorescence intensity) in WT14579 and Δ*cotE* sporangia at different sporulation stages (Fig. 3A, see also red boxes in Fig. 2A and Material and Methods). Before engulfment completion, the forespore and the mother cell compartments are located side by side (Fig. 1), and we therefore expected a FS/MC MTG fluorescence intensity ratio of 1 (Fig. 3A, red dashed line) towards the MCD pole of the forespore (Fig. 1). Just after engulfment completion, the forespore is surrounded by two membranes, in close proximity (Fig. 1) and the theoretical FS/MC MTG fluorescence intensity ratio is then equal to 2. In line with these predictions, the average FS/MC MTG fluorescence intensity ratio was 0.93±0.21 (median±standard deviation, n=153) and 0.99±0.23 (n=316) in WT14579 and Δ*cotE* sporangia respectively, at an intermediate stage of engulfment (Fig. 3A, “engulfing”, see red boxes in MTG channel of panel *a* of Fig. 2A). In sporangia with an engulfed forespore (red boxes in panel *d* of Fig. 2A), the FS/MC MTG fluorescence intensity ratio was 1.78±0.28 (n=211) and 1.81±0.29 (n=143) in WT14579 and Δ*cotE,* respectively (Fig. 3A). In sporangia with a TMR-star signal incompletely covering the surface of the refringent forespore (“incomplete”, red boxes in panel *e* of Fig. 2A), the FS/MC MTG fluorescence intensity ratio locally increased to 3.50±1.30 (n=92) and to 6.05±1.91 (n=131) in WT14579 and Δ*cotE* sporangia, respectively (Fig. 3A). These observations suggest that, in addition to the two membranes that delineate the forespore contour, the MTG dye associates with an additional structure on the forespore surface, giving a brighter signal in sporangia of refringent forespores. Finally, the FS/MC MTG fluorescence intensity ratio decreased to 2.08±0.56 (n=156) when the TMR-star homogeneously covered the forespore surface of WT14579 sporangia (“complete”, red boxes in panel *e* of Fig. 2A), only slightly higher than the ratio obtained in sporangia with no TMR-star signal on the surface of engulfed forespores (1.78±0.28) (Fig. 3A). In contrast, the ratio of FS/MC MTG fluorescence intensity was similar in Δ*cotE* sporangia of refringent forespores showing incomplete TMR-star labelling (6.05±1.91) and in sporangia with a TMR-star fully covering the forespore surface (5.37±2.04, n=44). Our quantification of the fluorescence signals suggests that, once fully assembled, the coat becomes less permeable to MTG in WT sporangia. However, this was not observed in Δ*cotE* sporangia, suggesting that coat properties are altered, even if the coat seems fully assembled on the forespore surface.

**Figure 3.**
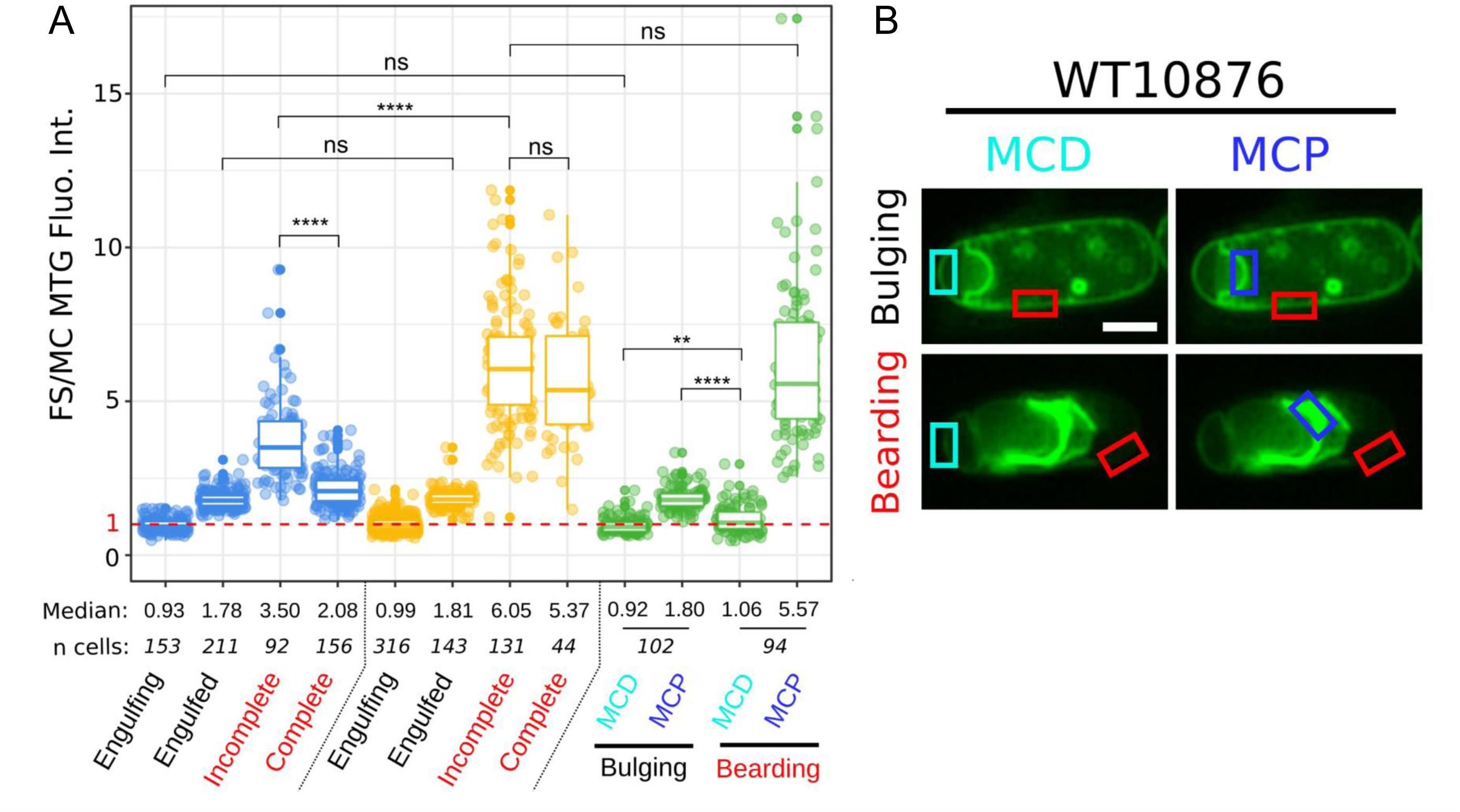
Quantification of MTG signal in sporangia at different sporulation stages. (**A**) Boxplots showing quantification of MTG fluorescent signals ratio intensity between the forespore (FS) and the mother cell (MC) membranes in WT ATCC14579 (blue boxes), Δ*cotE* (orange boxes) and in engulfment-defective (green boxes) sporangia in *B. cereus* ATCC 10876 sporulating at 20°C. The indicated sporulation stages refer to the Fig. 2A/B for WT14579 and Δ*cotE,* respectively. For engulfment-defective sporangia, quantification was performed towards the MCP and the MCD regions. (**B**) Illustrations of the regions used for recording the MTG fluorescence intensity in representative engulfment-defective cells. Cells are from at least two independent experiments and fluorescence intensity was recorded in pseudo-widefield images (see Materials and methods). Dashed red line represents a ratio of FS/MC MTG signal intensity=1. Each dot represents a different sporangia and n, the number of sporangia used for the quantification. The median values are indicated. The non-parametric Mann-Whitney U test was used, ns; not significant; **, p < 0.005; ****, p < 0.00005.

### The coat is partly assembled in engulfment-defective *B. cereus* cells

We previously reported that sporulation of *B. cereus* ATCC 10876 (WT10876) at 20°C leads to abnormal engulfment phenotypes in a variable proportion of the sporulating cells^15^. To determine whether formation of coat material depends on engulfment completion in *B. cereus,* we analyzed coat assembly in engulfment-defective WT10876 cells. We first confirmed, using a dual labeling of cell membranes with FM4-64 and MTG^48, 49^, that engulfment was indeed aborted in a fraction of the WT10876 sporulating cells at 20°C (Supplementary Fig. S1A, cyan arrows). Additionally, using a forespore fluorescent reporter (P*spoIIQ*-YFP), we observed bulging of the forespore towards the mother cell cytosol (Supplementary Fig. S1B, cyan arrows), a phenotype associated with impaired septal degradation of peptidoglycan^50–54^. In agreement with these observations, SIM revealed that the developing forespore pushes across the asymmetric septum and forms a bulge into the mother cell cytoplasm at hour 24 (Fig. 4A panels *a-c*). At hour 48, sporangia with a bulging phenotype never became refringent (Fig. 4A, panels *d-e* and Supplementary Fig. S1A, red asterisk), in contrast to sporangia with a normally engulfed forespore (Fig. 4A panel *g* and Supplementary Fig. S1A, yellow asterisks). Interestingly, TMR-star signals were detected in non-refringent-bulging sporangia (Fig. 4A, panels *d* and *e*, cyan arrows) and co-localized with bright MTG signals (curved and straight fragments; yellow and red arrows, respectively), suggesting the presence of coat material. Furthermore, the shape and the localization of TMR-star and MTG signals was strikingly similar to the particular mislocalization of coat fragments, also referred as “coat bearding”, previously observed in engulfment-defective *C. difficile* cells^35^. Finally, at hour 72, corresponding to the time of spore release (Fig. 4A, panel *f*, yellow asterisks), rare bulging sporangia were still present and we mainly detected abundant curved fragments (cyan arrows) sprinkled in the media, indicating that bulged sporangia ultimately lysed.

**Figure 4.**
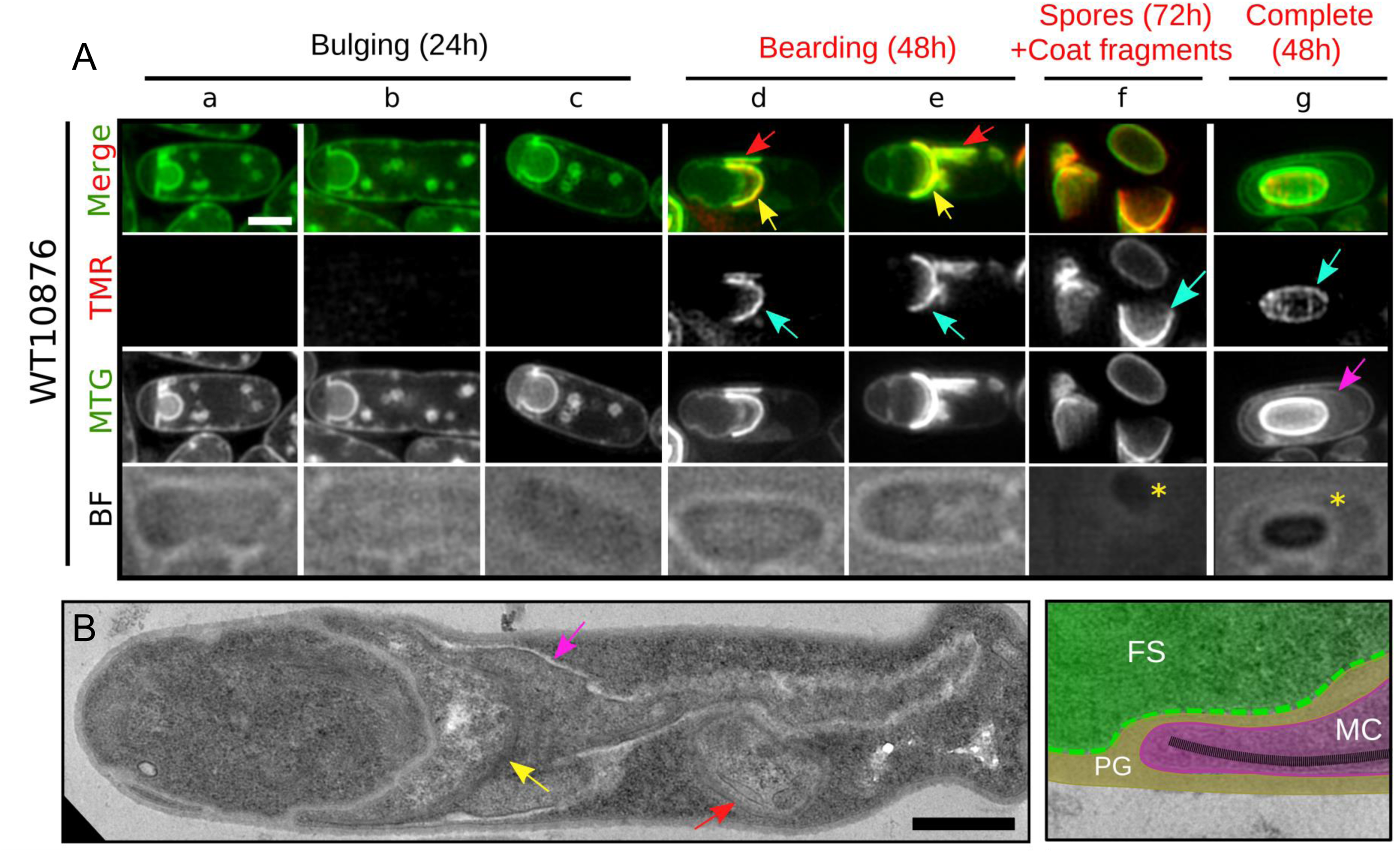
Fluorescence SIM and TEM analysis reveal a coat bearding phenotype in engulfment-defective *B. cereus* cells. WT10876 cells sporulating at 20°C in SMB were imaged (**A**) with SIM and BF illumination modes after labeling with TMR-star and MTG or (**B**) by TEM after thin sectioning. (**A**) Typical sporangia showing an engulfment-defective phenotype detected at hour 24 and not presenting TMR-star or MTG bright signals (“bulging”, panels *a-c*) or at hour 48 and showing TMR-star and MTG signals (“bearding”, panels *d-e*) are shown. Sporangia with an engulfed refringent forespore detected at hour 48 (panel *g*) and spores together with lysis fragments observed at hour 72 (panel *f*) are also shown for comparison. Cyan arrows indicate TMR-star signal, yellow arrows show curved MTG bright signal and red arrows point to MTG bright signal drawing a straight line in the medial focal plan. Pink arrow points to the exosporium. Scale bar is 1 μm. (**B**) WT10876 cells sporulating at 20°C, collected at hour 48 showing a blocked engulfment observed with TEM. Yellow arrow points to the large bearding coat fragment and red arrow points to the smaller fragments of coat disseminated in the MC cytoplasm. Pink arrow points at exosporium material. Right panel shows an inset of the junction between the FS (in green) and the MC (in magenta) compartments separated by peptidoglycan (PG) material (brown). Polymerized coat fragments (black dashed line) are present until the leading edge of the MC membranes. Scale bar represents 0.6 μm.

We performed a quantitative analysis of the FS/MC MTG fluorescence intensity ratio in WT10876 engulfment-defective sporangia (Fig. 3A and B, see also Materials and methods), showing TMR-star/MTG bright signals (“bearding”) or not (“bulging”). We performed the quantification for both the MCD and the mother cell proximal pole regions (MCP, here the forespore bulge) as the FS/MC MTG fluorescence intensity ratio should differ in these two regions of a partially engulfed forespore (See light and dark blue boxes in Fig. 3B). In the MCD region of bulging sporangia, the FS/MC MTG fluorescence intensity ratio was 0.92±0.25 (n=77) (Fig. 3A and B), confirming that only one membrane is present in this forespore region, while in the MCP region the ratio was 1.80±0.37 (n=77) (Fig. 3A and B), as in engulfed WT14579 forespores with no TMR-star signal (Fig. 3A). In sporangia showing a bearding phenotype, the FS/MC MTG fluorescence intensity ratio was 1.05±0.44 (n=166) at the MCD pole and 5.57±2.67 (n=166) at the MCP (Fig. 3A and B). This latest value is similar to that measured for Δ*cotE* sporangia with a refringent forespore (Fig. 3A). We concluded that MTG appears associated to a similar structure on the surface of Δ*cotE* forespores and in the front region of partly engulfed forespore of WT10876 sporangia.

In parallel, we imaged by TEM sporulating engulfment-defective cells collected at hour 48, at the time we detected MTG bright/TMR-star signals (Fig. 4B). According to our SIM observations, we clearly observed cells with an incomplete engulfment and coat material present towards the leading edge of the mother cell engulfing membrane (Fig. 4B, yellow arrow, see also inset in the right panel), together with a smaller amount of coat material inside the mother cell cytoplasm (Fig. 4B, red arrows). In addition, we clearly observed abundant exosporium-like material closely associated with the coat material (Fig. 4B, pink arrows). MTG signals corresponding to exosporium were not observed by SIM in bearding sporangia (Fig. 4A, panels *d* and *e*), likely hidden by the bright MTG signal bound to the coat material very close to the exosporium. Detection of abundant exosporium material in the cytoplasm of the mother cell indicates that extension of the exosporium from the cap, usually observed concomitantly with coat deposition^18, 23, 24^, also occurs in engulfment-defective *B. cereus* cells. In agreement with this, we observed partial encasement of the forespore bulge by CotE-SNAP and formation of a continuous CotE-SNAP signal in apparent engulfment-defective cells with MTG bright signals (Supplementary Fig. S2, see also Materials and methods). Altogether, these results confirm our ability to assess faithfully by SIM the state of coat development in live sporangia.

### The TMR-star signal on the coat is σ^K^-dependent

A previous study reported the absence of coat in Δ*sigK* sporangia of a *B. cereus sensu lato* group strain (*B. thuringiensis* 407, WT407)^55^. Thus, if our TMR-star labelling faithfully reports the electron-dense coat material localization described by TEM, no TMR-star labelling should be observed in Δ*sigK* sporangia. We imaged WT407 and Δ*sigK* sporangia after dual labeling with TMR-star and MTG, using conventional epifluorescence and phase contrast microscopy (Fig. 5A-D, upper panels). Here we focused our analysis on TMR-star signals since MTG poorly distinguished coat once completely assembled in a WT condition (Fig. 3A, blue boxplots, compare also panels *d* and *g* in MTG channel of Fig. 2A). The characteristic ovoid TMR-star signal was observed covering the surface of most of WT407 refringent forespores at hour 24 (Fig. 5A, “No fusion”, cyan arrows, see also the linescan profile in Fig. 5B), while it was undetected in Δ*sigK* sporangia (Fig. 5C, see also the linescan profile in Fig. 5D). This result shows that detection of TMR-star signal on the forespore surface is σ^K^-dependent.

**Figure 5.**
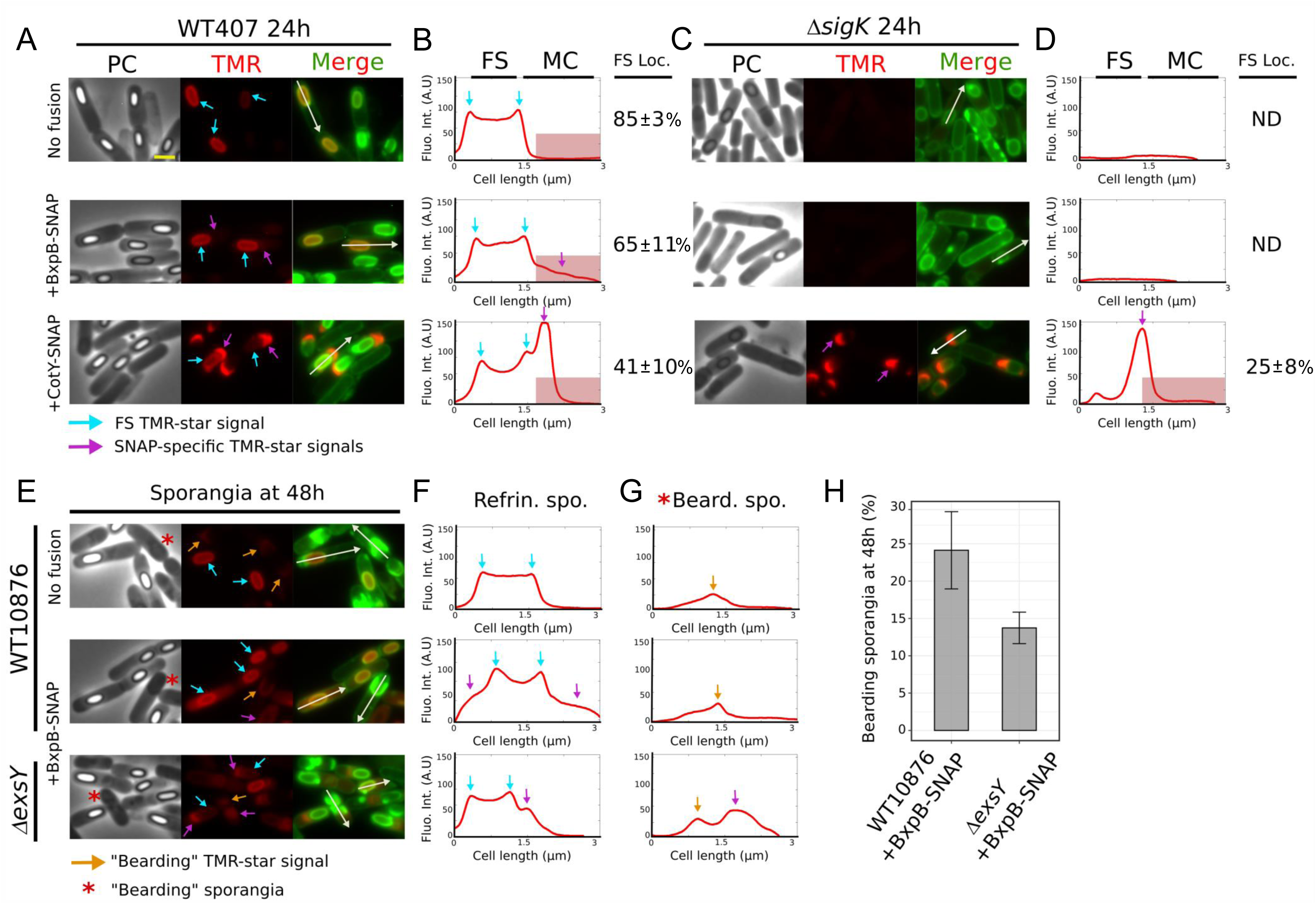
Detection of TMR-star signal on the forespore surface and specific BxpB-SNAP-TMR-star signal both depend on σ^K^. (**A-B**) WT407 or (**C-D**) Δ*sigK* cells sporulating at 20°C and expressing the indicated SNAP fusion were collected at hour 24, labeled with TMR-star (red) and MTG (green) dyes and imaged by conventional fluorescence microscopy and phase contrast (PC) microscopy. (**E-G**) Sporangia of WT10876 without SNAP fusion (“No fusion”), with BxpB-SNAP, and Δ*exsY* with BxpB-SNAP were collected after hour 48 and analyzed with conventional fluorescence microscopy and phase contrast (PC) microscopy after dual labeling with TMR-star and MTG. Red asterisks indicate sporangia with an apparent “bearding” phenotype. Orange arrows indicate TMR-star signal observed in engulfment-defective sporangia. Cyan arrows point to TMR-star binding on the surface of refringent forespore (FS) surface. Pink arrows point at the specific signal from TMR-star binding on indicated SNAP-fusion. White arrows indicate linescan used to quantify TMR-star fluorescence intensity along the indicated cell; the intensity line profile are shown in B, D, F and G. Red box highlights TMR-star signal localized in the mother cell (MC) cytoplasm and the mean fraction of the fluorescence intensity recorded in the FS is indicated (n = 25 cells; SD; standard deviation). A. U, Arbitrary Unit. ND, not determined. Scale bar in panel A represents 1 µm. Merged images represent TMR-star and MTG channels superimposed. (**H**) Percentage of “bearding” sporangia detected at hour 48 in the indicated strain (See Materials and Methods). Error bar represents SD of three biological replicates.

### A SNAP fusion allows visualization of BxpB, a late exosporium protein

In *B. cereus,* SNAP-fusions were particularly efficient to localize early assembling exosporium proteins such as CotE^15^, while a GFP fusion was not^13^. In addition, an attempt to localize a fusion of GFP to BxpB, a σ^K^-dependent late assembling exosporium protein^56^, failed in *B. cereus,* while equivalent fusions were successfully visualized in *B. anthracis* and *B. megaterium* ^22, 57^. To check whether i) BxpB can be localized using a SNAP-fusion, ii) BxpB-SNAP-TMR-star signal can be distinguished from binding of TMR-star on coat fragments and iii) σ^K^ is active in engulfmentdefective cells, we monitored by conventional fluorescence microscopy the detection of BxpB-SNAP in WT407 (Fig. 5A and B, middle panels), in a congenic Δ*sigK* mutant (Fig. 5C and D, middle panels), in WT10876 (Fig. 5E-G, middle panels) and in a congenic Δ*exsY* (Fig. 5E-G, lower panels) during sporulation at 20°C. In the absence of ExsY, exosporium extension does not occur and only the exosporium MCP cap with sometimes a thinner MCD cap, both formed by CotY, are assembled^15, 20, 25^. Importantly, in Δ*exsY* strains of *B. cereus* and *B. anthracis,* the exosporium MCP cap is less stably attached to the spore surface^15, 19, 20, 25^. Hence, in engulfmentdefective cells of a Δ*exsY* strain, we expected to reduce the spatial proximity between exosporium and coat material to maximize our chance to distinguish eventual BxpB-SNAP-TMR-star signals from TMR-star-coat signals.

In *B. anthracis,* BxpB-mCherry and –GFP fusions are firstly detected at low level in the mother cell cytoplasm and ultimately localize as a ring around the forespore^16, 22^. Accordingly, in addition to a characteristic ovoid TMR-star signal on the surface of refringent forespores and likely reflecting TMR-star binding to the coat (Fig. 5A and B, E and F, middle panels, cyan arrows), we observed a weak signal in the mother cell of WT407 and WT10876+BxpB-SNAP (Fig. 5A and B, E and F, middle panels, pink arrows and see red boxes in the linescan profiles). This signal was not observed in the absence of BxpB-SNAP (Fig. 5A and B, E and F, upper panels, “No fusion”). Later at hour 42, BxpB-SNAP displayed an exosporium localization (Supplementary Fig. S3) while no TMR-star signal was detected in Δ*sigK*+BxpB-SNAP sporangia (Fig. 5C and D, middle panels, see also Supplementary Fig. S3). In contrast, the cap localization specific of the early assembling CotY-SNAP was observed in WT407 and Δ*sigK* cells (Fig. 5A-D, lower panels, pink arrows and the linescans). Hence, together those results show that, despite the binding of TMR-star to the coat observed upon σ^K^ activation, a SNAP fusion allows the successful detection of a late assembling exosporium protein.

A weak TMR-star signal was observed in engulfment-defective WT10876+BxpB-SNAP cells (Fig. 5E and G, middle panels, red asterisks), however this signal was not specific from BxpB-SNAP since it was similar to the one observed in absence of the SNAP fusion (Fig. 5E and G, upper panels, orange arrows). In Δ*exsY* sporangia with a refringent forespore, in addition to a TMR-star signal on forespore surface (Fig. 5E and F, lower panels, cyan arrows), we observed a clear TMR-star signal specific of BxpB-SNAP drawing a cap-like structure towards the mother cell cytoplasm (Fig. 5E and F, lower panels, pink arrows). Bearding phenotypes were observed in Δ*exsY* sporangia but less frequently than in WT10876 sporangia (14±4% versus 24±9%, Fig. 5H). Importantly, similar cap-like localizations were also observed in engulfment-defective Δ*exsY* cells, suggesting binding of TMR-star to the BxpB-SNAP fusion (Fig. 5E and G, red asterisks in bottom panels, pink arrows and the linescan). However, apparent MTG bright signals indicating presence of coat material are often detected closely associated to the cap-like TMR-star signal in the mother cell cytoplasm. Thus, based on these diffraction limited images, we cannot fully rule out the possibility that the coat material is more spread in the mother cell cytoplasm of Δ*exsY* engulfment defective cells.

### BxpB-SNAP is specifically detected in engulfment-defective sporangia

To determine whether a specific TMR-star signal of BxpB-SNAP can be distinguished from TMR-star binding to coat material in engulfmentdefective Δ*exsY* cells, we designed a quantitative co-localization analysis of TMR-star and MTG signals (Fig. 6, see Materials and methods). We reasoned that signal originating from TMR-star bound on coat fragments should co-localize with MTG brighter signals, while a specific BxpB-SNAP-TMR-star signal should not. We performed this co-localization analysis on high-resolution lattice-SIM images of spores, of exosporium cap and of engulfment-defective Δ*exsY* cells showing brighter MTG signals (“bearding”) (Fig. 6A and B). We used

**Figure 6.**
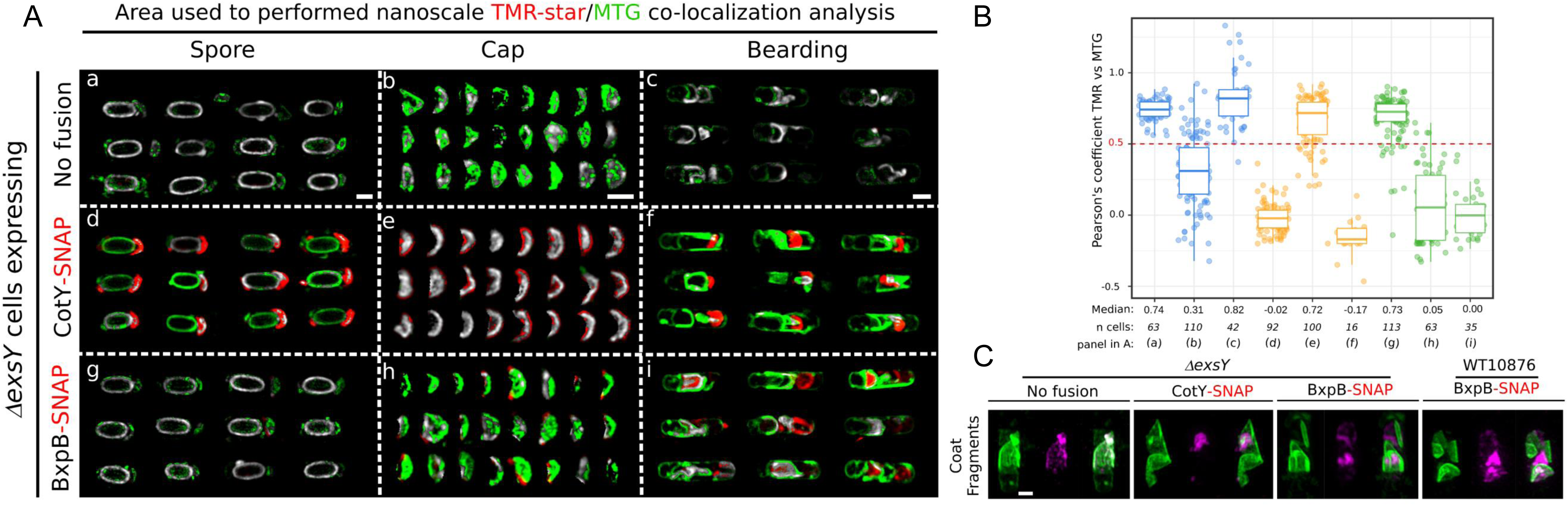
BxpB-SNAP-TMR-star signal is specifically detected in sporangia showing a bearding pheno-type. (**A-B**) Co-localization analysis for TMR-star and MTG signals in Δ*exsY* spores, in exosporium cap or in bearding sporangia expressing the indicated SNAP fusion. Sporulation was performed in SMB at 20°C, collected at hour 72, labeled with TMR-Star and MTG dyes and imaged with lattice-SIM. (**A**) Representative colocalization maps generated by the co-localization threshold plugin on Fiji. Grey color indicates co-localization of MTG and TMR-star signals, green color indicates a preferential MTG signal enrichment and red color indicates predominant TMR-star signals. Scale bars represent 0.8 μm. (**B**) Boxplots showing quantification of Pearson coefficient for TMR-star and MTG signals in Δ*exsY* without fusion (blue boxes), with CotY-SNAP (orange boxes) or with BxpB-SNAP (green boxes). A dashed red line indicates a positive correlation (r>0.5). Median values are indicated. Each dot represents a different sporangia and n, the number of sporangia used for quantification. The median values are indicated. (**C**) Z-projections of coat fragments released after lysis of bearding cells. TMR-star signals are in magenta and MTG signals represented in green. Scale bar represents 0.8 μm.

Δ*exsY* cells without SNAP-fusion (No fusion) to monitor the co-localization of TMR-star and MTG signals due to their binding on coat fragments and Δ*exsY*+CotY-SNAP cells, as a positive control of a specific exosporium TMR-star binding.

In *exsY* spores without SNAP fusion, TMR-star and MTG signals largely co-localized in the central region of the spore (grey in panel a, Fig. 6A), with a Pearson’s correlation coefficient “r” of 0.74±0.07 (n=63) (Fig. 6B). In contrast, the TMR-star signal did not co-localize with the MTG signal around the forespore in Δ*exsY*+CotY-SNAP spores (Fig. 6A panel *d*, r=0.02±0.10, n=92) and appeared instead preferentially associated with the cap(s) (Fig. 6A and 6B). Accordingly, when looking exclusively at the exosporium cap, the TMR-star signal co-localized with the MTG for Δ*exsY*+CotY-SNAP (panel *e*, r=0.67±0.19, n=100), while a random distribution of the correlation coefficient (r=0.31±0.25, n=110) is observed in Δ*exsY* spores without fusion (panel *b*). Importantly, the TMR-star signal did not co-localize with the MTG in Δ*exsY*+CotY-SNAP bearding sporangia (panel *f*, r=-0.16±0.14, n=16), in contrast to Δ*exsY* sporangia without fusion (panel *c*, r=0.83±0.21, n=42). Hence, the CotY-SNAP-TMR-star signal can be specifically distinguished from the TMR-star binding to the coat material in Δ*exsY* engulfment-defective sporangia.

The analysis of Δ*exsY*+BxpB-SNAP cells shows that the TMR-star signal co-localized with MTG signal in spores (r=0.69±0.16, n=113, panel *g* in Fig. 6A, and 6B) but not in the exosporium cap of spores (panel *h*, r=0.07±0.26, n=63). Interestingly, in this latest condition, the TMR-star appears often associated with the margin of the exosporium cap, a particular pattern of localization which was recently reported by an immunofluorescence analysis of BxpB in *B. anthracis* Δ*exsY* mutant spores^19^. In sporangia showing the “bearding” phenotype, we observed a strong TMR-star signal which did not co-localize with MTG (panel *i*, r=-0.01±0.19, n=35). We concluded that the TMR-star signal observed in Δ*exsY*+BxpB-SNAP engulfmentdefective sporangia that does not colocalize with MTG, reveals the specific binding of TMR-star on BxpBSNAP.

We also acquired Z-stack images of the coat fragments released by the lysis of bearding cells using lattice-SIM (Fig. 6C). On maximum projections of Z-stack images, we clearly recognized specific TMR-star signal of CotY-and BxpB-SNAP fusions (in magenta) in the 3D space drawn by coat fragments (in green) labeled by the MTG (Fig. 6C). In contrast, in Δ*exsY* without fusion, the TMR-star signal is only viewed superimposed on MTG signals. This analysis also showed a specific TMR-star signal due to BxpB-SNAP, associated to the fragments released after the lysis of bearding WT10876 sporangia (Fig. 6C). In this latest condition, the BxpB-SNAP-TMR-star signal appears closely associated to the curved coat fragments. Taken together, both the nanoscale co-localization analysis and super-resolved Z-maximum projection images demonstrate the presence of a TMR-star signal specific of BxpB-SNAP in engulfment-defective *B. cereus* cells, suggesting that σ^K^ is active in these cells.

## Discussion

Determination of the state of coat and exosporium development in sporangia or in spores is an unavoidable step in developmental studies of *B. cereus*. To date, this was largely performed by TEM. We recently showed that MTG labeling allows to directly observe the basal layer of the exosporium cap after engulfment completion and the following steps of exosporium formation using SIM^15^. Here, we demonstrated that both TMR-star and MTG specifically stain the coat of *B. cereus*, and that SIM reconstructed signals can be used to assess the precise stages of coat development. Thus, it is now possible using these fluorescence markers to determine the precise stage of both exosporium and coat assembly in living cells using fluorescence microscopy, without the need for genetic manipulations. Since DNA recombination is poorly effective and the genetic toolbox is limited for *B. cereus* species compared to other *Bacilli*^11, 12^, we believe that our non-genetic fluorescence live imaging approach could be particularly efficient to characterize mechanisms of proteinaceous spore surface layers assembly in new *B. cereus* isolates or to assess the effect of destruction treatments at the cell level. Notably, we showed that binding of TMR-star to the coat gives a characteristic ovoid signal around the forespore in WT14579, WT10876 and WT407 strains. Two of those strains are *B. cereus sensu stricto* species while WT407 is a *B. thuringiensis* strain. Hence, we assume that our results are valid for other species of the *B. cereus sensu lato* group and notably for *B. anthracis* strains.

Although TMR-star and MTG both bind to the coat, the detected signals differ due to the ability of MTG to bind also to the celĺs membranes and exosporium, while TMR-star only binds to the coat. Interestingly, the signal originating from TMR-star binding the coat could still be detected in some strains expressing SNAP-fusions. Notably, in cells expressing BxpB-SNAP, binding of TMR-star to the coat firstly represents most of the TMR-star signal observed (Fig. 5A and B, middle panels). In a WT strain expressing CotE-, CotY-or ExsY-SNAP fusions, such TMR-star association with the forespore surface was not distinguishable at any stage of the sporulation^15^. Encasement by those morphogenetic proteins is completed soon after engulfment completion and before the forespore becomes refringent, and the strong signal of these fusions in vicinity of the forespore surface likely masks the weak TMR-star signal coming from binding to the coat. In agreement with this statement, the TMR-star association with the forespore surface can be observed when localization of CotY-SNAP is impaired, as observed in Δ*cotE*, Δ*exsY*^15^ and in WT407 strains (Fig. 5A-B, lower panels). Hence, our studies indicate that TMR-star cannot be used as a coat marker in strains expressing abundant early assembling SNAP-tag exosporium proteins. In such strains, bright MTG signals reconstructed by SIM and observed at intermediate stages of coat deposition or in cells showing coat assembly defects can be used as spatial and temporal markers of coat development.

In this study, we also find that the coat appears less permeable to MTG when completely covering the forespore surface, as judged by a complete ovoid TMR-star signal observed in such cells and in free spores. We also noticed that the coat material accessibility to MTG is higher in Δ*cotE* sporangia, including those with a forespore fully covered by a TMR-star signal, suggesting that coat permeability is affected among the whole population of Δ*cotE* sporangia. Hence, the high sensitivity to lyzozyme reported for Δ*cotE* spores formed at 20°C^47^ may not only be due to the partly decoated spores, as observed by TEM. A recent report indicates that CotG, a CotE-controlled protein of *B. subtilis* (ExsB is a homologue in *B. cereus*^58, 59^), and a central determinant of outer coat patterning^60–63^, mediates spore permeability^64^. Thus, it is tempting to suggest that a defect in ExsB assembly may explain the apparent increase accessibility of the coat in *B. cereus* Δ*cotE* cells formed at suboptimal temperature. Our observations show that the quantification of the MTG signals is a promising tool to evaluate the structuration state and likely the permeability of the coat directly at the single cell level.

We applied our super-resolved SIM-based fluorescent analysis to characterize the proteinaceous spore surface layers development in a heterogeneous population, observed when sporulation *B. cereus* ATCC 10876 cells occurs at 20°C. We confirmed a complete arrest of the engulfment process in a variable part of the cells, with: i) a dual labeling of cell’s membrane with MTG and FM4-64, ii) a TEM analysis and iii) a quantification of MTG fluorescence signals. Engulfment defects observed were characterized by a bulging of the forespore towards the mother cell cytoplasm. These bulging cells never became refringent and ultimately lysed, indicating a failure in the forespore development. Importantly, despite an incomplete engulfment, we characterized multiple features normally observed only in engulfed cells; i) synthesis and accumulation of coat material close to the forespore bulge, creating a coat bearding phenotype, ii) partial encasement by CotE-SNAP, iii) exosporium extension and iv) synthesis of BxpB-SNAP fusion. Except the encasement by CotESNAP, the other features are governed by the late mother-cell specific σ^K^ factor. Altogether, our results suggest that σ^K^ is active in engulfment-defective WT10876 cells.

Strikingly, the coat bearding phenotype reported here in engulfment-defective *B. cereus* cells, is reminiscent of the one described in engulfment-defective *C. difficile* sporangia^35^. Bearding phenotype in *C. difficile* likely originates from the absence of an inhibitory σ^K^ pro-sequence^29^. In *B. cereus* genomes, a σ^K^ prosequence and associated protease are present^65^. However, in most of the *B. cereus* species, an interrupting *skin* element and the associated SpoIVCA recombinase are absent^66, 67^. Thus, *B. cereus* and *C. difficile* had evolved different strategies leading to a less tight regulation of σ^K^ activation. Importantly, since modulation in σ^K^ activation control has been only demonstrated in pathogenic species, one can suggests that σ^K^ activity in engulfment-defective cells may confer an advantage in the context of an infection. An appealing possibility is that the formation and release of partially structured materials of the proteinaceous spore surface may be the way to expose to the media/the host, internal components otherwise never exposed. Such exposition may create new interactions with the immune system, to dupe it and/or to generate a particular matrix for the normally assembled spores.

## Materials and methods

### Bacterial strains, plasmids and general methods

Strains and plasmids used in this study are listed in Table 1. The various fluorescent fusions are carried by the low copy pHT304-18 plasmid and described in Table 1. The *bxpB-SNAP* gene fusion was synthesized by ATUM (www.atum.bio, Newark, CA). Briefly, synthetic inserts corresponding to the *bxpB* gene (bc_1221) and the 169 bp sequence upstream of the *bxpB* start codon of *B. cereus* ATCC 14579, followed by a GCAGCTGCT linker and the SNAP sequence were inserted into the pHT304-18 plasmid, using *Sal* I and *EcoR* I restriction sites, giving rise to the pHT304-BxpB-SNAP (Table 1). All plasmids were first introduced into *E. coli* DH5α and clones were confirmed by PCR and/or DNA sequencing. Plasmids were then transferred to *E. coli* SCS110; the resulting unmethylated plasmids were then transferred to the *B. cereus/B. thuringiensis* strains by electroporation. LB agar plates or LB broth with orbital shaking (200 rpm) were used for routine growth at 37°C. When needed, liquid cultures or plates were supplemented with the following antibiotics at the indicated concentrations: ampicillin (Amp) at 100mg·ml^−1^ for *E. coli* cultures, spectinomycin (Spc) at 275mg·ml^−1^, kanamycin (Kan) at 200mg·ml^−1^ and erythromycin (Erm) at 5mg·ml^−1^ for *B. cereus* and *B. thuringiensis* cultures. Sporulation was induced by resuspension in liquid SMB medium at 20°C with orbital shaking at 180 rpm, as previously described^15^.

**Table 1.**
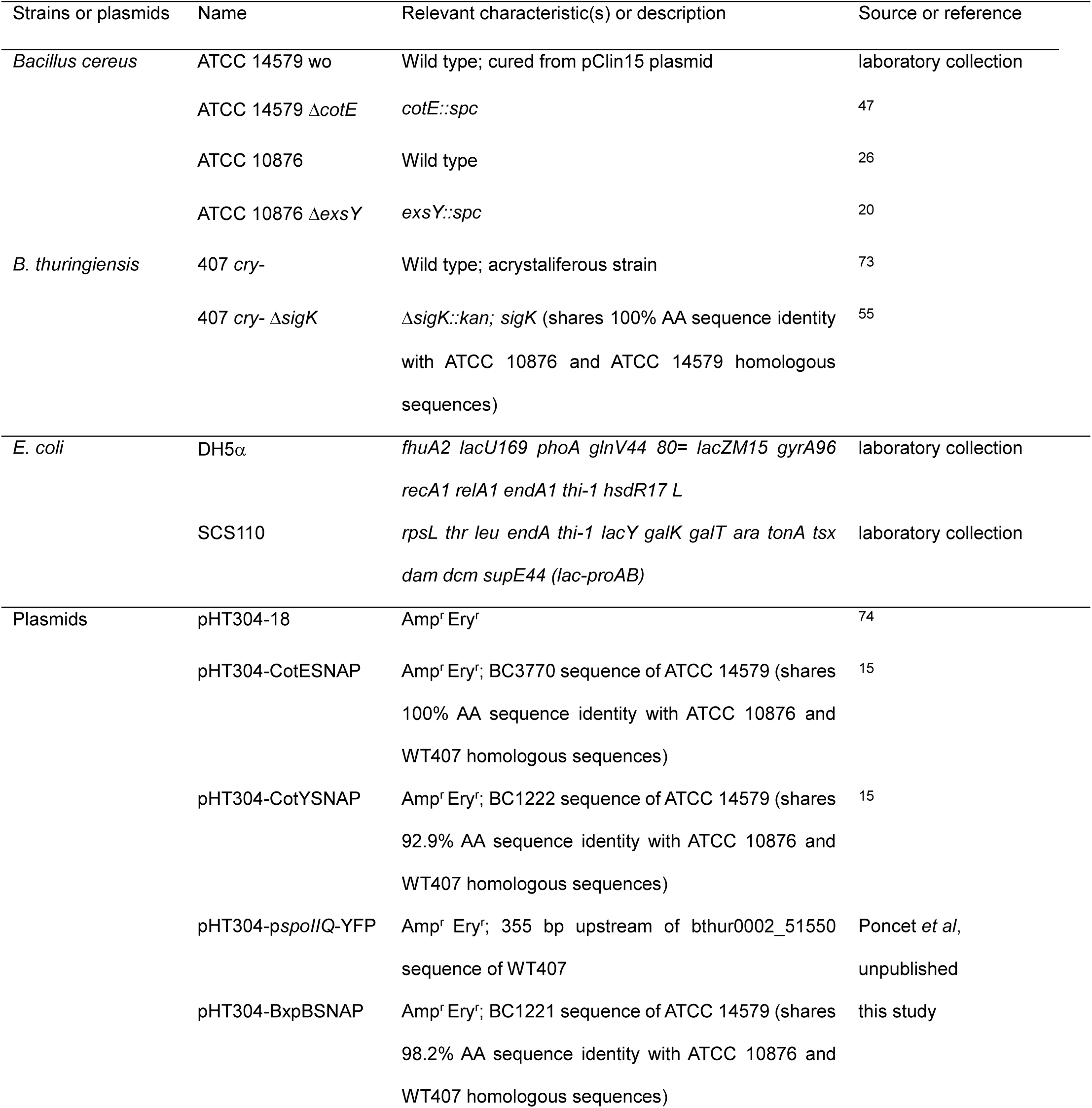
Strains and plasmids used in this study.

### SNAP labeling and fluorescence microscopy

Samples (1mL) were withdrawn from cultures in SMB at selected times during sporulation. Cells were collected by centrifugation (10,000 x g for 3 min), resuspended in 200 µL of phosphate saline buffer (PBS) and labeled by incubation with SNAP-cell TMR-Star (New England Biolabs) for 30 min at 37°C in the dark at a final concentration of 250 nM. This TMR-star-probed suspension was centrifuged (12,000 x g, 1 min), washed with 1 mL of PBS, suspended again in 1mL of PBS and labeled with Mitotracker Green (MTG, Thermofischer) and/or FM4-64 Fx at a final concentration of 1 µg/mL or in 100 nM MV405 for simultaneous visualization of YFP protein (Supplementary Fig. S3) for 1 min at room temperature. Cells were then washed three times in PBS and suspended in 50µL to 200 µL PBS, depending on the concentration of sporulating cells/spores. For diffraction-limited as well as super-resolved microscopy, 3 µL of the labeled cells suspension were applied onto 1.7% agarose in PBS-coated glass slides. All experiments were done at RT. Samples were imaged with a BX-61 (Olympus), Elyra PS.1 or Elyra 7 AxioObserver (Zeiss) microscope.

### Microscopy acquisition settings

Samples were observed with an epifluorescence microscope (BX-61; Olympus) equipped with an Olympus UPlanFL N 100x/1.30 Oil Microscope Objective, an Orca Flash 4.0 LT camera (Hamamatsu) and illuminated by a fluorescence LED lamp (PE300 WHT-365). Images were acquired using the CellSens Olympus software, for TMR-star acquisitions exposure time was 500ms for all images and 31ms for MTG acquisitions. Final pixel size was 65 nm for raw image. Super Resolution Structured Illumination Microscopy (SIM)^68^ images were acquired using an Elyra PS.1 Microscope (Zeiss) equipped with a Plan-Apochromat 63x/1.4 oil DIC M27 objective and a Pco. edge 5.5 camera, using 488 nm (100mW) or 561 nm (100 mW) laser lines, at 15% and 10% of total potency for 488 and 561 nm lasers respectively. Final pixel size was 64.5 nm for raw image. The grid periods used were 28 mm or 34 mm for acquisitions with the 488 nm or 561 nm lasers respectively. For each SIM acquisition, the corresponding grating was shifted and rotated five times, giving 25 acquired frames. Final SIM images were reconstructed using the ZEN software (black edition, 2012, version8, 1, 0, 484; Zeiss), using synthetic, channel-specific Optical Transfer Functions (OTFs), baseline cut and noise filter settings ranging from −7 to −8. Lattice SIM imaging was performed on a Zeiss Elyra 7 AxioObserver inverted microscope equipped with 488 nm (100 mW), 561 nm (100 mW), and 642 nm (150 mW) laser lines, at 20% of maximal output power, a sCMOS camera (PCO Edge edge 4.2), a 63×/NA 1.4 objective (Zeiss, Plan-Apochromat 63x/1.4 Oil DIC M27) and an additional lens (1.6x) in the detection pathway. Final pixel size was 64.5 nm for raw image. The fluorescence emission was separated from the laser excitation by a dichroic beamsplitter (405/488/561/641) and filtered (dual band with two bandpass 495-550 and 570-620) before detection. Image acquisition was controlled by the Zen Software (Zeiss, black edition). The Lattice SIM approach is based on stands on an illumination of point pattern^69, 70^ and laterally shifted patterns on the sample (called phase images). The raw images were composed of 15 phase images per plane per channel, and acquired with a Z-distance of 0.100 μm. The grids were chosen to be optimal for both laser lines and modulation contrast (27.5 and 32 µm grids for the 488 and 561 laser lines respectively). The lattice SIM reconstruction was performed with Zen software with automatic settings, baseline cut and sharpness settings ranging from 7 to 9. Negative values resulting from SIM processing (due to the sharpness filter) were clipped by setting them to zero. The intensities of final reconstructed SIM images were normalized to the maximum dynamic range to be displayed with optimal brightness and contrast. For all exposure time was 20 ms.

### Image analysis

All image analysis were performed and micrographs processed using Fiji software (ImageJ, NIH). Brightness and contrast of representative cell images were adapted in Fiji and figures were compiled using Inkscape 0.92.5. (https://inkscape.org). At least three different microscopic fields were analyzed for every condition.

### (i) Quantification of MTG signal ratio intensity between the forespore and the mother cell

The sporulation stages were determined with brightfield (BF) and MTG channels. First, on SIM reconstructed images, one polygonal region of interest (ROI) was adjusted using the Fiji polygon tool on the forespore (FS) membrane and a second ROI was positioned on the mother cell (MC) membrane of the same sporangia. The polygonal ROI were transferred on pseudo-widefield images and the mean intensity in the two ROI were recorded and an individual FS/MC was computed for every sporangium. This method was repeated for the indicated numbers of sporangia. At least two biological replicates were used per condition. We used Δ*cotE*+CotY-SNAP SIM images previously acquired in the same condition^15^ as a second replicate for quantification of FS/MC MTG ratio in Δ*cotE* cells. Since assembly of CotY-SNAP is blocked in *ΔcotE* cells, a minimal impact of CotY-SNAP synthesis on coat deposition is attempted. To increase the size of our statistical sampling of engulfment-defected cells, we used sporangia in different ATCC 10876 backgrounds, showing engulfment defects that we used in the course of the present study. Hence results presented in Fig. 3A included ratio computed for WT10876, WT10876+CotE-SNAP, Δ*exsY*, Δ*exsY*+CotE-SNAP strains.

### (ii) Quantification of interspace size on SIM images

The distance separating the basal layer of the exosporium cap from the underlying outer forespore membrane (OFM) was calculated in MTG channel of SIM images using the Fiji function ‘points to distance’. Only sporangia with an apparent exosporium cap in the focal plan were used. The distance was calculated between the exosporium cap point at the maximum distance from the surface of the forespore and the center of the MCP pole of the forespore.

### (iii) Counting of sporangia and spores with defects of coat assembly

SIM images of sporangia of refringent spores (black in BF images) from two biological replicates of WT cells collected at hour 24 or Δ*cotE* and Δ*cotE* expressing CotY-SNAP cells, collected at hour 48 or spores collected at hour 72 were used for analysis. The Δ*cotE* cells presented a delay of sporulation and none of the engulfed sporangia observed at hour 24 was refringent or showed TMR-star/MTG brighter signals. Normal patterns of MTG brighter signals localization were defined according to previous TEM study^15, 18, 23^ and as presented in Fig. 1.

### (iv) Quantification of TMR-star signal in the forespore

Quantification was adapted from Delerue et al^71^. To determine the proportion of TMR-star signal associated with the forespore among the sporangia, a polygonal ROI was first fitted on the forespore region and a second ROI was positioned on the sporangia outline in the TMR-star channel of diffraction-limited images. For Δ*sigK* sporangia, since forespore did not become refringent, we adjusted ROI on engulfed cells with phase black/phase grey forespore. The total fluorescence was recorded in the forespore (F_FS_) and in the sporangium (F_sporangium_). The forespore signal was computed for individual sporangia as follow: (F_FS_-F_background_)x100/(F_sporangium_-F_background_). The mean fluorescence of the image signal background (F_background_) was adjusted for every microscopy field.

### (v) Quantification of sporangia showing a coat bearding phenotype among the sporulating population on diffraction-limited images

The proportion of sporangia showing a bearding phenotype at hour 48 among the global sporulating population was determined by identifying cells with a red TMR-star signal, an apparent forespore bulging as judged by MTG labelling and not appearing bright or grey in phase contrast microscopy. Biological triplicates were used for W10876+BxpB-SNAP and Δ*exsY*+BxpB-SNAP cells.

### (vi) Quantification of Pearson correlation coefficient on Lattice-SIM images

Co-localization of MTG with TMR-star signals on lattice-SIM reconstructed images (see “Microscopy acquisition settings section”) was assessed by the Colocalization Threshold plugin^72^ available on Fiji. The co-localization coefficient was recorded in different regions defined using the polygon tool of Fiji in MTG channel and encompassing, (i) the entire spores including the associated cap(s), (ii) only the biggest cap or (iii) the contour of the sporangia showing a bearding phenotype. Pearson’s correlation coefficient was calculated for individual regions, with a coefficient value of 1 representing a perfect positive linear correlation and a value of 0 indicating an absence of correlation. When no correlation was found by the function, a score of –0.2 was assigned.

### (vii) Quantification of TMR-star signal length

Quantification was performed on SIM reconstructed images (Supplementary Fig. S2C and D). Using Fiji program, a segmented line was fitted from one end to the other of the layer drawn by the TMR-star signal. For CotE-SNAP, additional, weakest layers were often observed. Quantification was only performed on the layer closest from the forespore bulge. The length of the line, used as a proxy of encasement by CotE-SNAP and CotY-SNAP was determined in the different sporangia classified using bright field and MTG fields. Sporangia showing an apparent forespore bulging, a FS/MC MTG signal ratio towards the MCD pole <1.2, were defined as “bulging” cells and those showing, in addition, MTG bright signals towards the bulge, *i.e.* a FS/MC MTG ratio >3, were defined as “bearding” cells.

### Transmission electron microscopy

Samples from SMB cultures were collected by centrifugation 48 h after the onset of sporulation, and the cells fixed and processed for thin sectioning TEM as described before^15^.

### Statistics

Statistical analyses were performed using Rstudio version 4.1.1 for PC. Data are represented with boxplots showing the interquartile range (25^th^ and 75^th^ percentile). The upper whisker extends from the hinge to the largest value no further than 1.5 x IQR from the hinge and the lower whisker extends from the hinge to the smallest value at most 1.5 x IQR of the hinge. All sporulation kinetics were replicated at least twice using conventional fluorescence microscopy before being imaged by SIM or lattice SIM. The number of cells analyzed is indicated in each figure.

## Data availability

The datasets used and/or analyzed during the current study are available from the corresponding author on reasonable request.

## Acknowledgments

We thank Pr Mariana Pinho for providing access to the SIM PS1 microscope, Pedro Matos for very efficient training on image acquisition and analysis software and Dr Cécile Morlot for fruitful discussions. We also thank Dr Leyla Slamti for the gift of WT407 and Δ*sigK* strains, Pr Anne Moir for Δ*exsY* and Δ*cotY* mutant strains and Dr Sandrine Poncet for pHT304-p*spoIIQ*-YFP plasmid.

## Author contributions

Conceptualization, A.L., V.B., A.O.H, F.C; methodology, A.L., S.C., C.B., M.S., I.B.; formal analysis, A.L., V.B., visualization, A.L.; writing original draft, A.L.; writing, review and editing, A.L., V.B., A.O.H, F.C, RCL; project administration and funding acquisition, V.B., supervision, V.B., F.C. All authors have read and approved the final manuscript.

## Additional information

Supplementary figures S1-S3

