## Supplementary material for "A new fluorescence-based approach for direct visualization of coat formation during sporulation in *Bacillus cereus*": Suplementary Figures S1 S2 S3

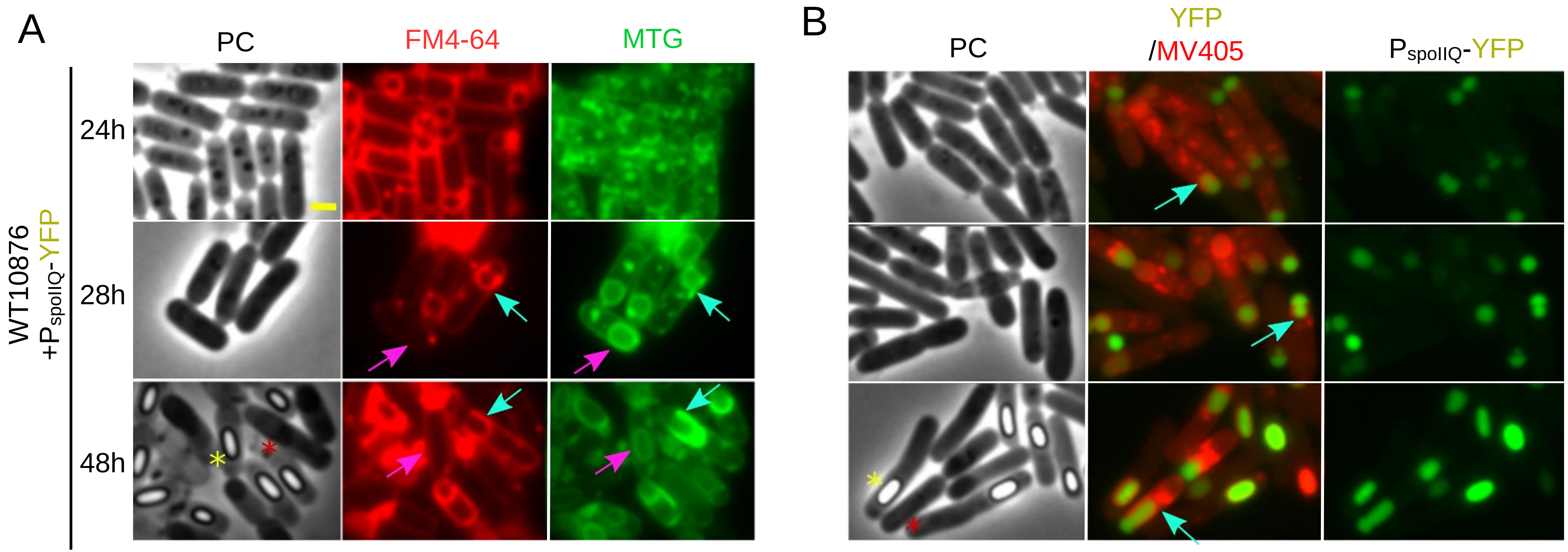


**Figure S1. Sporulation at 20°C leads to a forespore bulging which prevents engulfment completion in a subpopulation of sporulating WT10876 cells.** *B. cereus* WT10876 cells expressing a P*spoIIQ*-YFP transcriptional fusion as a marker of the forespore and sporulating at 20°C were collected at indicated times and imaged by phase contrast (PC) and fluorescence microscopy, after labeling with the membrane dyes FM4-64 (red) and MTG (green) (A) or MV405 (red) (B). After engulfment completion, MTG stains the forespore membranes (pink arrows) while the lipophilic FM4-64 did not (pink arrows). At hours 28 and 48, the forespore bulge is labelled by both MTG and FM4-64 showing an engulfment defect (cyan arrows). MV405 membrane dye (false colored red) was used to simultaneously visualized YFP protein (false colored green). The YFP is expressed under the *spoIIQ* promoter and confirms the forespore bulging phenotype (cyan arrows). Yellow asterisks indicate refringent forespore and red asterisks point to engulfment-defective cells. Scale bar in A represents 1 µm.

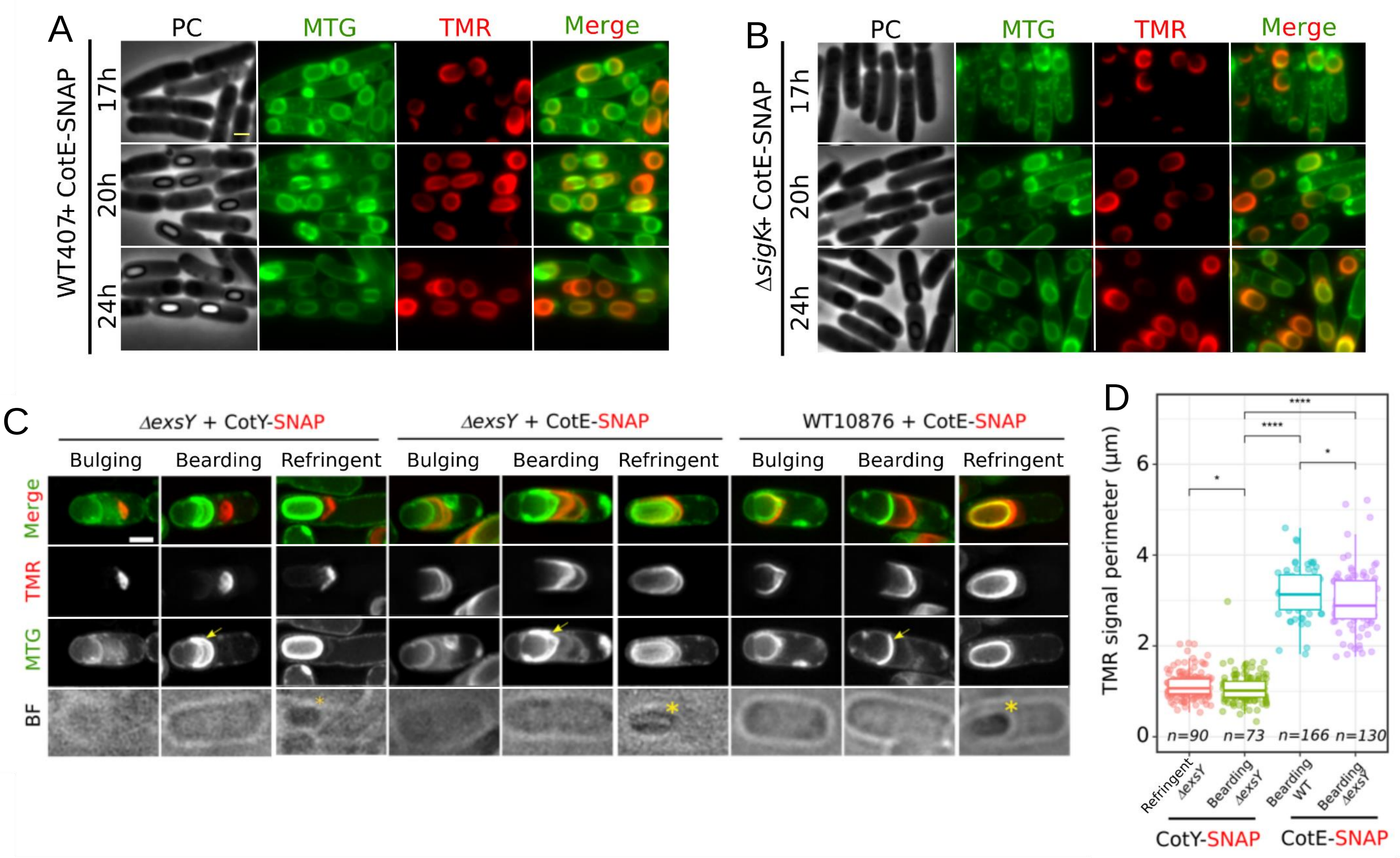


**Figure S2. The $\sigma^K$–independent CotE-SNAP encasement partly occurs in engulfment-defective sporangia.** WT407 (A) or congenic Δ*sigK* cells (B) both expressing CotE-SNAP and sporulating at 20°C were collected at indicated times, labeled with TMR-star (red) and MTG (green) dyes and imaged by conventional fluorescence and phase contrast (PC) microscopy. A complete encasement by CotE-SNAP is observed in Δ*sigK* and WT cells. (C-D) Δ*exsY* cells with CotY-SNAP or CotE-SNAP fusion or WT10876 cells with CotE-SNAP sporulating at 20°C were collected at hour 48 and labeled with the TMR-star and MTG dyes and imaged by SIM and BF microscopy. In engulfed Δ*exsY* cells ("refringent"), CotY-SNAP encasement is blocked while encasement by CotE-SNAP is completed, similarly to the WT (Refringent). In "bearding" cells, encasement by CotY-SNAP is also blocked, drawing a small cap, as observed in sporangia with a refringent forespore. In contrast, CotE-SNAP assembles as a large array in both bearding Δ*exsY* and in WT bearding cells, indicating that extension of the CotE-SNAP layer, usually observed only after engulfment completion, occurs in engulfment-defective cells. Yellow arrows point to MTG bright signals in bearding cells and yellow asterisks indicate refringent forespore. (D) Boxplots showing quantification of TMR-star signal perimeter used as a proxy of SNAP fusion encasement. Bulging and bearding cells were defined based on the FS/MC MTG signal ratio calculation, see Materials and methods. Each dot represents a sporangia. Scale bars in A and C represent 1µm. The non parametric Mann-Whitney U test was used, ns, non significant; *, $p < 0.05$, ****, $p < 0.00005$, n; numbers of sporangia used for quantification.

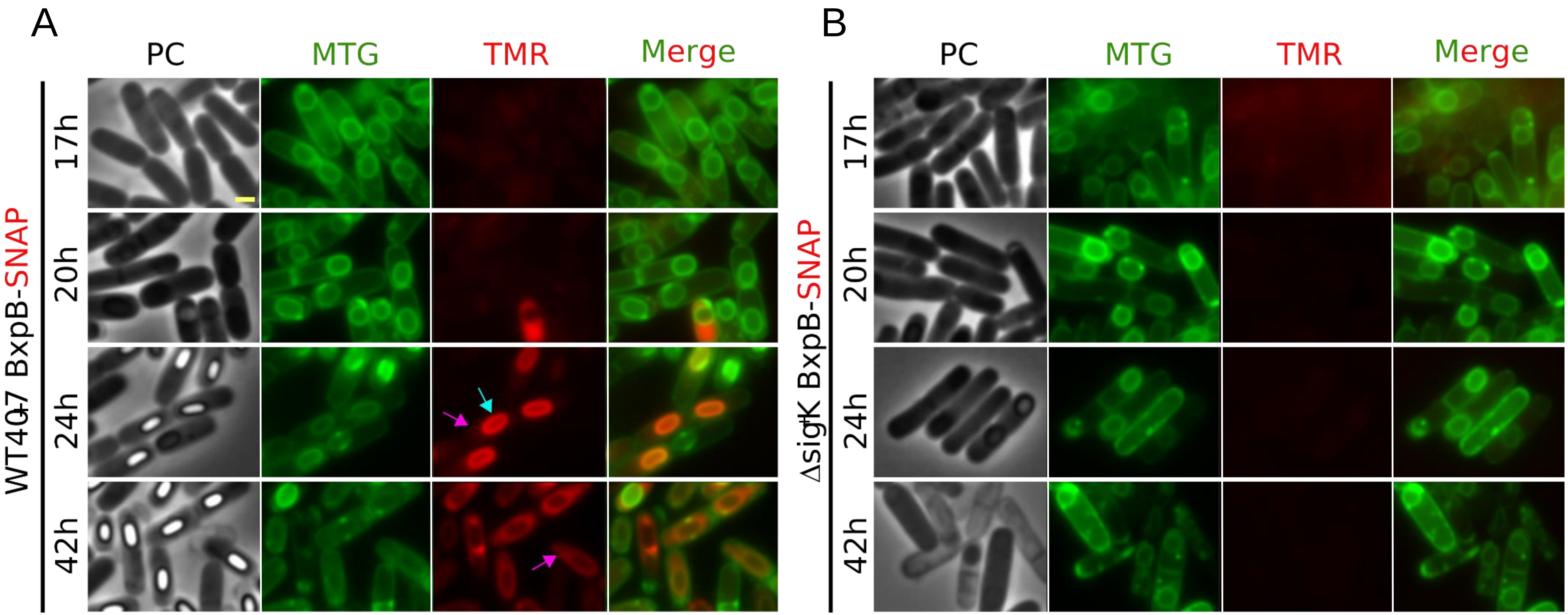


**Figure S3. Control of BxpB-SNAP assembly in WT407 and Δ*sigK* sporulating cells.** (A) WT407+BxpB-SNAP or (B) Δ*sigK*+BxpB-SNAP cells sporulating at 20°C collected at indicated times were labeled with TMR-star (red) and MTG (green) dyes and imaged by fluorescence and PC microscopy. At hour 24, the TMR-star signal coming from dye binding to the coat (cyan arrow) is stronger than the BxpB-SNAP-TMR-star signal (pink arrows). At hour 42, an exosporium-like localization of BxpB-SNAP-TMR-star is mainly observed in sporangia of refringent forespores, while the TMR-star signal due to binding on the coat cannot be distinguished. Scale bar in A represents 1 µm.
